# Infrared light stimulates the mitochondrial large-conductance calcium-activated potassium channel in guinea pig cardiomyocytes

**DOI:** 10.64898/2026.06.11.731586

**Authors:** Joanna Lewandowska, Piotr Bednarczyk, Barbara Kalenik, Bogusz Kulawiak, Antoni Wrzosek, Adam Szewczyk

**Affiliations:** Laboratory of Intracellular Ion Channels, Nencki Institute of Experimental Biology, Polish Academy of Sciences, Warsaw, Poland; Department of Physics and Biophysics, Institute of Biology, Warsaw University of Life Sciences, Warsaw, Poland

**Author notes:** Equal contribution (Joanna Lewandowska, Piotr Bednarczyk).

**Keywords:** cardiomyocytes, cytochrome c oxidase, mitochondria, potassium channels, infrared light

## Abstract

Mitochondrial potassium channels play an important role in regulating cellular metabolism, redox balance, and survival, particularly in excitable tissues such as the heart. Among them, the mitochondrial large-conductance calcium-activated potassium (mitoBK_Ca_) channel has been implicated in cardioprotection during ischemia–reperfusion injury. At the same time, growing evidence indicates that mitochondria act as light responsive organelles, with cytochrome c oxidase (COX) serving as a primary chromophore for red and near-infrared (NIR) light. In this study, we investigated whether 820 nm infrared light modulates mitoBK_Ca_ channel activity in mitochondria isolated from guinea pig cardiomyocytes. Using patch-clamp recordings of mitoplasts, we demonstrated that illumination at 820 nm NIR wavelength enhanced mitoBK_Ca_ channel activity in a redox-dependent manner. Our findings reveal a previously unrecognized mechanism linking NIR light modulation via COX to the regulation of cardiac mitoBK_Ca_ channels as a metabolic sensor. This study identifies the mitoBK_Ca_ channel as a novel effector of light-induced mitochondrial signaling and suggests that modulation of cardiac mitochondrial potassium transport by NIR light may contribute to cardioprotective effects. These results provide new insight into the integration of bioenergetic and photoregulatory processes in mitochondria and support the development of non-pharmacological strategies targeting mitochondrial function.

## INTRODUCTION

Mitochondria are not only central regulators of cellular metabolism but also integrate bioenergetic processes with redox signaling and cell survival pathways (Chandel, 2014; Galluzzi et al., 2012; Gorman et al., 2012; Murphy, 2009). In cardiac cells, mitochondrial function is particularly critical due to the high and continuous demand for ATP. Among the key elements controlling mitochondrial physiology in cardiomyocytes are ion channels located in the mitochondrial membrane, such as mitochondrial porin (VDAC, voltage-dependent anion channel) or calcium uniporter (Kwong et al., 2015; O’Rourke, 2007). These also include mitochondrial potassium (mitoK) channels present in the inner mitochondrial membrane, which regulate membrane potential, matrix volume, and reactive oxygen species (ROS) synthesis (Garlid et al., 2003; O’Rourke, 2004; Szabo & Szewczyk, 2023; Szabo & Zoratti, 2014). In the inner mitochondrial membrane in cardiac cells, a few different mitoK channels were identified (Augustynek et al., 2017).

One of the most widespread potassium channels in the inner mitochondrial membrane is the large-conductance calcium-activated potassium (mitoBK_Ca_) channel. It was first described using the electrophysiological patch-clamp technique in astrocytoma cells (Siemen et al., 1999). Discovery of the mitoBK_Ca_ in the inner mitochondrial membrane of cardiac cells demonstrated its important role in protecting the heart from ischemic injury (Xu et al., 2002). Using patch-clamp recordings, fluorescence-based K⁺ flux measurements, and biochemical analyses, the authors showed that this large-conductance channel is activated by Ca²⁺, contributes substantially to mitochondrial K⁺ uptake, and is distinct from the previously known mitoK_ATP_ channel (Xu et al., 2002). Pharmacological experiments reveal that activation of mitoBK_Ca_ enhances mitochondrial K⁺ influx, whereas its inhibition markedly suppresses it. Importantly, in isolated heart models, activation of mitoBK_Ca_ significantly reduces infarct size, the effect abolished by specific channel blockers, indicating a direct cytoprotective function. Mechanistically, the mitoBK_Ca_ channel is proposed to regulate mitochondrial ion homeostasis, volume, and bioenergetics, and to limit Ca²⁺ overload and oxidative stress during ischemia. Overall, the findings establish mitoBK_Ca_ channels as a key component of mitochondrial physiology and highlight mitochondrial K⁺ transport as a general mechanism underlying cardioprotection, with potential therapeutic implications (Szteyn & Singh, 2020).

Accumulating evidence indicates that mitoK channels are not isolated entities but interact functionally and structurally with mitochondrial proteins, including components of the respiratory chain (Kathiresan et al., 2009; Peng et al., 2014; Sokolowski et al., 2011). For example, mitochondrial ATP-sensitive potassium (mitoK_ATP_) channels have been proposed to interact with succinate dehydrogenase (Ardehali et al., 2004), while mitochondrial TASK-3 (mitoTASK-3) channels are also linked to respiratory chain components (Yao et al., 2017). Similarly, the mitochondrial Kv1.3 channel has been shown to associate with complex I (Peruzzo et al., 2020). In cardiac mitochondria, the β1 subunit of the mitochondrial large-conductance calcium-activated potassium (mitoBK_Ca_) channel interacts with cytochrome c oxidase (COX) subunit I (Ohya et al., 2005), and additional interactions between mitoBK_Ca_ channels and respiratory complexes have been reported in both cardiac (J. Zhang et al., 2017) and brain mitochondria (Singh et al., 2016).

This growing body of evidence suggests a direct functional coupling between the energy-generating system (the respiratory chain) and the energy-dissipating system (potassium channels), representing an important regulatory mechanism within mitochondria. In line with this concept, we previously demonstrated that mitoBK_Ca_ channel activity in glioblastoma U-87 MG cells is modulated by mitochondrial substrates and inhibitors acting through the respiratory chain (Bednarczyk et al., 2013). Together, these findings point to cytochrome c oxidase (COX) as a potential key regulator of mitoBK_Ca_ channel function.

In parallel, increasing attention has been directed toward photobiomodulation (PBM), defined as the use of low-intensity red or near-infrared (NIR) light (620–850 nm) to modulate cellular metabolism and survival pathways (Karu, 2010; Zhang et al., 2021). PBM has been shown to enhance mitochondrial respiration, ATP synthesis, and redox balance, leading to protective or stimulatory effects depending on the physiological context (Karu et al., 2004). Beneficial outcomes of PBM have been reported across a range of conditions, including neurodegenerative disorders, cardiovascular diseases, and wound healing, suggesting that mitochondria harbor intrinsic photoresponsive elements that drive adaptive metabolic responses (Abijo et al., 2023). However, the molecular mechanisms underlying light-induced mitochondrial regulation remain incompletely understood.

Cytochrome c oxidase (complex IV, COX) is widely considered the primary mitochondrial chromophore responsible for the absorption of red and near-infrared light, particularly at wavelengths such as 620, 680, 760, and 820 nm. This property is attributed to its metal centers, including copper and heme groups, which undergo reversible redox transitions upon photon absorption (Karu, 2010; Karu et al., 2004). Light-induced activation of COX can enhance enzymatic turnover, stimulate electron transport, and improve oxygen utilization, ultimately increasing ATP synthesis under both physiological and stress conditions (Hamblin, 2018). Interestingly, specific wavelengths can differentially regulate mitochondrial function: while 808 nm light typically stimulates COX activity, 750 nm and 950 nm wavelengths exert an inhibitory effect (Sanderson et al., 2018). This bidirectional control is critical in the context of ischemia-reperfusion injury, where the application of inhibitory wavelengths (750/950 nm) prevents mitochondrial hyperpolarization and the subsequent catastrophic release of reactive oxygen species (ROS) (Pham et al., 2025). Furthermore, Hüttemann’s work emphasizes that this non-invasive “metabolic pacing” can protect highly aerobic tissues, such as the brain and heart, during acute stress events (Wider et al., 2023). Collectively, these findings have shifted the field of photobiomodulation from simple metabolic enhancement to a sophisticated, wavelength-specific regulatory tool for cellular bioenergetics.

Despite substantial progress in understanding both photobiomodulation and mitochondrial ion channel regulation, the mechanism underlying the light-induced modulation of mitoBK_Ca_ channel activity remains unknown and whether such regulation depends on the redox state of the respiratory chain and the illumination wavelength. In particular, no comprehensive study has addressed whether NIR light within the COX absorption spectrum can influence mitoBK_Ca_ channel gating. Recent studies indicate NIR light stimulates mitoBK_Ca_ channel activity, with cytochrome c oxidase acting as a key photoacceptor (Bednarczyk et al., 2026).

In this study, we investigated whether 820 nm near-infrared light within the effective absorption range of cytochrome c oxidase modulates mitoBK_Ca_ channel activity in mitochondria isolated from guinea pig cardiomyocytes. Using patch-clamp electrophysiology, we demonstrate that mitoBK_Ca_ channel activity is regulated by 820 nm light in a redox-dependent manner, revealing a novel link between photobiomodulation and cardiac mitochondria potassium channel function. This effect may contribute to the cardioprotective effects induced by 820 nm NIR light.

## MATERIAL AND METHODS

### Chemicals

Acrylamid, Bis–Tris, Coomassie blue G–Brillant Blau G250, glycerin, potassium hydroxide, and tricine were obtained from Carl Roth GmbH + Co., Germany. Digitonin and n-dodecyl-ß-D-maltoside were obtained from SERVA, Germany. Ammonium persulfate, potassium chloride, and tris(hydroxymethyl)aminomethane were purchased from BioShop, Canada. PBS–Dulbecco’s phosphate buffered saline w/o magnesium and w/o calcium was from Biowest, France. Protease inhibitor cocktail tablets complete EDTA-free were from Roche, Germany. RNase-free DNase set was from Qiagen AG, Germany. Tween-20 was purchased from Bio-Rad, USA. All other chemicals were from Sigma-Aldrich, USA.

### Cardiomyocytes isolation form guinea pig heart

All investigations involving animals conformed to the Guidelines for the Care and Use of Laboratory Animals published by the US National Institutes of Health, and the experimental procedures used in this study were approved by the local Animal Research Committee of the Nencki Institute of Experimental Biology. Adult male Dunkin Hartley SPF guinea pigs (HsdDhl:DH) were euthanized by cervical dislocation. Cardiomyocytes were isolated with Langendorf perfusion system (Frankenreiter et al., 2017) with minimal changes described in (Lewandowska et al., 2026). After thoracotomy, the hearts were harvested and placed in a warm Perfusion Buffer containing: 113 mM NaCl, 25 mM KCl, 0.1 mM KH_2_PO_4_, 0.6 mM Na_2_HPO_4_, 1.2 mM MgSO_4_, 0.03 mM Phenol red, 11.6 mM NaHCO_3_, 10.1 mM KHCO_3_, 10 mM HEPES, 30 mM Taurine, 500 mM 2,3-Butanedione monoxime, 1 g/l glucose, pH 7.4. After removal of residual blood from the coronary circulation with Perfusion buffer, its modification containing liberase and trypsin was used. The aorta and atrias were trimmed off. Next, the resulting part of the heart was cut into small pieces. Digestion was stopped by adding 2.5 ml of Stop Solution containing Perfusion buffer with 10% FBS. The resulting suspension was filtered through a 400 µm mesh. The filtrate was centrifuged at 20 g for 5 min at 25 °C. Quantity and quality of isolated cardiomyocytes were assessed with a light microscope (Fig. 1A) and a Neubauer chamber. The isolation procedure was considered successful when a minimum yield of 6.0 × 10⁵ rod-shaped cardiomyocytes was achieved.

**Figure 1.**
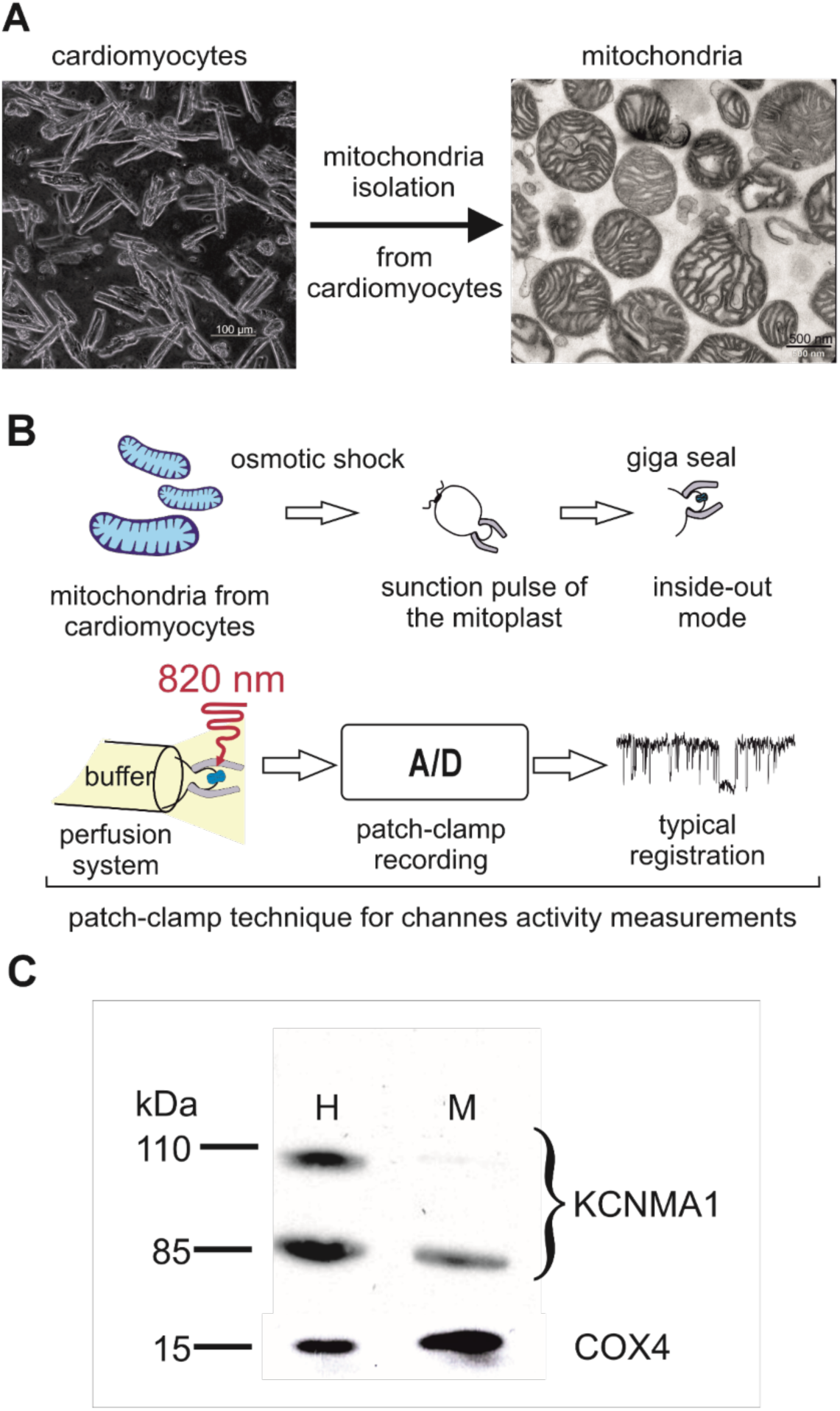
Microscopic characterization of isolated cardiomyocytes and mitochondria, experimental setup for patch-clamp recordings, and protein detection by Western blot analysis. (A) Representative light microscopy image of isolated parental cardiomyocytes and electron microscopy image of mitochondria obtained from these cells, confirming successful mitochondrial isolation and preservation of mitochondrial structure. (B) Schematic presentation of single-channel activity measurements performed using the patch-clamp technique. Isolated mitochondria were subjected to osmotic shock to obtain mitoplasts, followed by giga-seal formation and recordings in the inside-out configuration. Channel activity was recorded using a patch-clamp amplifier and analyzed after analog-to-digital conversion. (C) Western blot analysis demonstrating the presence of the α-subunit of the large-conductance calcium-activated potassium channel (BK_Ca_) and the mitochondrial marker cytochrome c oxidase subunit 4 isoform 1 (COX4) in cardiomyocyte homogenates (H) and isolated mitochondrial fractions (M) derived from guinea pig cardiomyocytes.

### Mitochondria isolation from cardiomyocytes

Mitochondria were isolated using a rapid, efficient method reported in detail by Lewandowska et al. (Lewandowska et al., 2026). Briefly, cardiomyocytes were subjected to gentle disruption in a buffer composed of 250 mM sucrose, 5 mM HEPES, and 1 mM EGTA (pH 7.2), supplemented with 1% bovine serum albumin (BSA) ussing cell strainers, providing functional mitochondria almost exclusively from cardiomyocytes. The quality of isolated mitochondria was confirmed in a few isolations with an electron microscope (Fig. 1A) and other methods described in (Lewandowska et al., 2026).

### Electron microscopy imaging

All TEM imaging samples were prepared according to the same protocol. If not indicated otherwise, each step was followed by centrifugation 2 min, 6000 rpm. Briefly, pellets were washed 6 times with 0.1 M cacodylate buffer. Then, samples were treated with 2% osmium tetroxide in 0.1 M cacodylate buffer for 60 min in the dark. After 3 x washing with water, the samples were treated with 2% uranyl acetate in water for 40 min in the dark. After 3 x washing with water, samples were dehydrated for 10 min in each concentration of acetone dilutions: 25%; 50%; 75%; 90%; 96% and finally 15 min in 100% acetone. Then samples were impregnated by resin - acetone:epon resin 1:1, 1h, and acetone:epon resin 1:2, 1h; epon resin 1h; epon resin o/n. During resin impregnation steps, centrifugation at 2 min, 7000 rpm was used. Then, the pellets were polymerized at 60 °C for 3 days. Next, samples were cut on ultrathin sections (70 nm) using an ultramicrotome, contrasted with 2% lead citrate, and viewed under a microscope.

Samples for TEM imaging and electron microscopy images were performed by the Laboratory of Electron Microscopy (Nencki Institute of Experimental Biology PAS) with high-resolution transmission electron microscope JEM 1400 (JEOL Co., Japan, 2008) with 11 Megapixel TEM Morada G2 camera (EMSIS GmbH, Germany).

### Calcium ions reintroduction to cardiomyocytes isolation buffer

Because the cardiac perfusion buffer was entirely Ca²⁺-free and the digestion buffers contained only low Ca²⁺ concentrations (approximately 50 µM), a gradual reintroduction of calcium was required prior to experiments assessing cardiomyocytes viability under control and hypoxic conditions. Immediate elevation of extracellular Ca²⁺ to 1.8 mM resulted in acute cellular damage; therefore, the final concentration was achieved through stepwise supplementation of Ca²⁺ to the isolation buffer. Specifically, Ca²⁺ concentration was increased sequentially to 200 µM, 600 µM, 1.0 mM, and ultimately 1.8 mM. Each increment was separated by a 10 min equilibration period, and the procedure was performed at 37 °C. Following the reintroduction of calcium ions, the buffer was replaced with DMEM supplemented with 10% FBS.

### Cardiomyocytes viability assay

Experiments were performed as previously described (Liu et al., 1996). Isolated cardiomyocytes were centrifuged at 20×g for 3 minutes. The cell pellet was suspended in culture buffer and divided into groups: control and those irradiated with 820 nm light.

### Cardiomyocyte irradiation with 820 nm light was performed in an incubator equipped with an

820 nm LED chamber for 15 minutes at 37 °C. Cells from both groups were then incubated for 1.5 hours and then divided into four groups: control and irradiated cells intended for incubation in normoxia, and control and irradiated cells intended for incubation in hypoxia, in equal 1.5 ml aliquots into Eppendorf tubes. Half of the control cells and cells preconditioned with 820 nm light were hypoxied by removing excess medium, leaving only 100 μl of cell suspension, which was carefully overlaid with a 400 μl layer of mineral oil. All cells were incubated in a thermoblock at 37 °C for 1 hour. Cell viability was assessed by trypan blue exclusion, counting stained and unstained cells under a microscope.

### Patch-clamp experiments with cardiac mitoplasts

Patch-clamp experiments were performed as previously described (Kampa et al., 2021; Walewska et al., 2022). In brief, mitoplasts were isolated from cardiomyocytes’ mitochondria by placing them in a hypotonic solution (5 mM HEPES, 100 μM CaCl_2_, pH 7.2) for approximately 2 minutes to induce swelling and rupture of the outer membrane (Figure 1B). A hypertonic solution (750 mM KCl, 30 mM HEPES, 100 μM CaCl_2_, pH 7.2) was added to restore isotonicity. The final bath isotonic solution contained 150 mM KCl, 10 mM HEPES, and 100 μM CaCl_2_ at pH 7.2. Similarly, the patch-clamp pipette was filled with an isotonic solution. Mitoplasts are easily identifiable by their size, rounded shape, transparency, and the presence of a “cap”, distinguishing them from cellular debris in the preparation. For all experiments, an isotonic solution containing 100 μM CaCl_2_ was used as the control. A low-calcium isotonic solution (1 μM CaCl_2_) contained 150 mM KCl, 10 mM HEPES, 1 mM EGTA, and 0.752 mM CaCl_2_ at pH 7.2. Channel modulators were added as dilutions in the isotonic solutions containing either 100 μM CaCl_2_ or 1 μM CaCl_2_. A perfusion system was used to apply these substances, consisting of a holder with a custom-made glass tube, a peristaltic pump, and Teflon tubing. Mitoplasts at the tip of the measuring pipette were transferred into the openings of a multibarrel “sewer pipe” system, where their outer faces were rinsed with the test solutions. All experiments were conducted in patch-clamp inside-out mode. The voltages reported correspond to those applied to the inside of the patch-clamp pipette, with positive potentials indicating the physiological polarization of the inner mitochondrial membrane (outside positive).

The electrical connection was established using Ag/AgCl electrodes and an agar salt bridge (3 M KCl) as the ground electrode. The current was recorded with a patch-clamp amplifier (Axopatch 200B, Molecular Devices, USA). Pipettes, made of borosilicate glass, had a resistance of 10–20 MΩ and were pulled using a Narishige PC-10 puller. The current was low-pass filtered at 1 kHz and sampled at 100 kHz. Single-channel recordings were analyzed, with channel conductance calculated from the current–voltage relationship. The probability of channel opening (Po) was determined using Clampfit software. The single-channel search mode was used for analysis. Results are presented as mean values ± standard deviation (S.D.). In figures showing single-channel recordings, “-” indicates the channel’s closed state.

### NIR illumination protocol

The illumination set-up contains: 150 W halogen lamp, MSH-150 monochromator systems, and glass fiber bundles (380 – 1600 nm) (Quantum Design). The beam was positioned on the patch-clamp pipette at 550 nm (green) (Figure 1B). The final power density at 820 nm was 335 μW/cm^2^. The light power densities were measured using a Microscope Slide Power Meter Sensor Head S170C (Thorlabs).

### SDS-PAGE and immunoblotting

Cardiomyocytes were seeded in 6-well plates, washed with cold PBS, and detached from the wells. Cells were collected by centrifugation and lysed in RIPA buffer supplemented with protease inhibitors. After incubation on ice, the lysates were clarified by centrifugation at 4 °C. The resulting supernatants were collected and stored at −20 °C for further analysis. Aliquots were taken for protein concentration determination prior to electrophoretic analysis. Equal amounts of whole-cell lysates solubilized in Laemmli buffer (Bio-Rad) were separated by 10% Tris–tricine SDS-PAGE and subsequently transferred onto polyvinylidene difluoride (PVDF) membranes (Bio-Rad). After transfer, membranes were blocked with 10% non-fat dry milk prepared in Tris-buffered saline containing Tween 20 and incubated with primary antibodies: anti-BK_Ca_ (Alomone Labs, APC-107, 1:500) and anti-COX4 (Cell Signalling, 1:200, 4844S). Immunodetection was performed using HRP-conjugated secondary antibodies against rabbit or mouse IgG (GE Healthcare and Thermo Fisher Scientific) followed by visualization with enhanced chemiluminescence reagents (GE Healthcare). Molecular weight estimation was carried out using the PageRuler Prestained Protein Ladder (Thermo Fisher Scientific).

### Statistical analysis

All experiments were performed with at least three or more independent biological replicates to ensure reproducibility. Results are presented as mean ± S.D., as calculated using Prism 4 (GraphPad Software Inc.). One-way ANOVA was employed to analyze the experimental data. P-values were considered significant as follows: *p ≤ 0.05, **p ≤ 0.01, and ***p ≤ 0.001.

## RESULTS

To investigate whether near-infrared (NIR) light modulates mitoBK_Ca_ channel activity in cardiac mitochondria, we designed a sequential experimental strategy combining structural, biochemical, electrophysiological, and functional approaches. First, we validated the quality of isolated cardiomyocytes and mitochondrial preparations to ensure reliable single-channel recordings. Next, we characterized the biophysical and pharmacological properties of the detected mitochondrial potassium channel to confirm its identity as mitoBK_Ca_. Having established a functional experimental model, we then examined whether 820 nm infrared light regulates channel activity under defined redox conditions associated with cytochrome c oxidase function. Finally, we assessed whether NIR illumination exerts a cytoprotective effect on isolated cardiomyocytes exposed to hypoxic stress, linking the observed electrophysiological effects to cellular survival.

### Quality of the cardiac mitochondria preparation

The quality of mitochondrial isolation is crucial for detecting single-channel activity in the inner mitochondrial membrane. In Figure 1A, we show a light microscopy image of parental guinea pig cardiomyocyte morphology and an electron microscopy image confirming the successful isolation of intact mitochondria from these cells. The isolated mitochondria display characteristic ultrastructural features, indicating good preservation of mitochondrial integrity after the isolation procedure. In Figure 1B, the experimental workflow for measuring single-channel activity using the patch-clamp technique is illustrated. Isolated mitochondria were subjected to osmotic shock to obtain mitoplasts; a giga seal was then formed, and recording was performed in the inside-out configuration. Channel activity was measured using the patch-clamp system, and signals were digitized through an analog-to-digital converter. Where indicated, the recordings were performed under 820 nm illumination, enabling analysis of channel behavior under experimental conditions. In the next step, Western blot analysis of proteins detected in cardiomyocyte homogenates (H) and isolated mitochondria (M) was performed. The α-subunit of the large-conductance calcium-activated potassium channel, was identified in both homogenates and mitochondrial fractions, indicating the presence of this potassium channel in cardiac mitochondria. In addition, COX4, a mitochondrial marker protein, was detected in both samples, confirming the successful enrichment of mitochondrial material in the isolated fraction. Together, these results validate the mitochondrial isolation procedure and demonstrate the presence of BK_Ca_ channels in mitochondria derived from guinea pig cardiomyocytes.

### Characterization of the channel present in the inner mitochondrial membrane

The biophysical and pharmacological properties of the detected channel were evaluated using the patch-clamp technique with classical modulators: calcium and paxilline (Pax). In Figure 2A, representative single-channel recordings obtained in a symmetric 150/150 mM KCl solution containing 100 µM Ca²⁺ demonstrate clear voltage-dependent channel activity. Channel openings increased with membrane depolarization, with the highest activity observed at positive membrane potentials (+40 to +60 mV). At negative voltages, channel activity was markedly reduced. These recordings confirm the presence of functional voltage-sensitive mitoBK_Ca_ channels in the inner mitochondrial membrane. Additionally, the current–voltage (I–V) relationship was derived from single-channel recordings (Figure 2C). The relationship was linear across the tested voltage range, indicating typical potassium-selective conductance under symmetric ionic conditions. The calculated single-channel conductance was 127 ± 2 pS, consistent with that of BK_Ca_ channels in cardiac cells. Parallelly, the voltage dependence of the open probability (Po) of the mitoBK_Ca_ channel in the presence of 100 µM Ca²⁺ was illustrated (Figure 2D). Channel open probability increased progressively with increasing membrane potential over the range from −80 to +120 mV, reaching maximal activation at positive voltages. These findings demonstrate strong voltage sensitivity of the channel. Next, in figure 2B, representative current traces recorded at +40 mV show the effects of calcium and pharmacological inhibition with paxilline (Pax) on channel activity. Under control conditions, spontaneous channel openings were observed. Reducing calcium concentration to 1 µM markedly decreased channel activity, whereas increasing it to 100 µM enhanced channel opening frequency. Application of 1 µM paxilline, a selective BK_Ca_ channel inhibitor, almost completely abolished channel activity. Finally, the voltage dependence of the open probability (Po) of the mitoBK_Ca_ channel in the presence of 100 µM Ca²⁺ was confirmed. The following set of results represents the electrophysiological characterization of the mitoBK_Ca_ channel recorded in mitoplasts isolated from guinea pig cardiomyocytes in the presence of NS11021, a classical potassium channel opener. In panel A, representative recordings obtained at +40 mV show channel activity under sequential experimental conditions. Under control conditions, the mitoBK_Ca_ channel displayed a relatively high level of activity. The addition of 1 µM Ca²⁺ markedly reduced channel openings, indicating inhibition of channel activity at this calcium concentration. Subsequent application of 3 µM NS11021 in the presence of 1 µM Ca²⁺ restored channel activity, demonstrating that NS11021 effectively activates mitoBK_Ca_ channels even under inhibitory calcium conditions. Increasing the calcium concentration to 100 µM further enhanced channel opening frequency and duration, consistent with strong calcium-dependent activation of mitoBK_Ca_ channels. Finally, application of 1 µM paxilline almost completely abolished channel activity, confirming the pharmacological identity of the recorded currents as BK_Ca_-mediated. The quantitative analysis of channel open probability is summarized in Figure 3B. Under control conditions, the open probability remained high, whereas exposure to 1 µM Ca²⁺ significantly decreased it to nearly zero. Application of 1 µM Ca²⁺ and 3 µM NS11021 partially restored channel opening. The highest variability and strong activation were observed in the presence of 100 µM Ca²⁺, where open probability reached substantially elevated values. In contrast, treatment with 1 µM paxilline reduced the open probability to almost undetectable levels. Together, these results demonstrate that mitoBK_Ca_ channel activity is strongly regulated by calcium concentration and can be pharmacologically enhanced by NS11021 while being effectively inhibited by paxilline.

**Figure 2.**
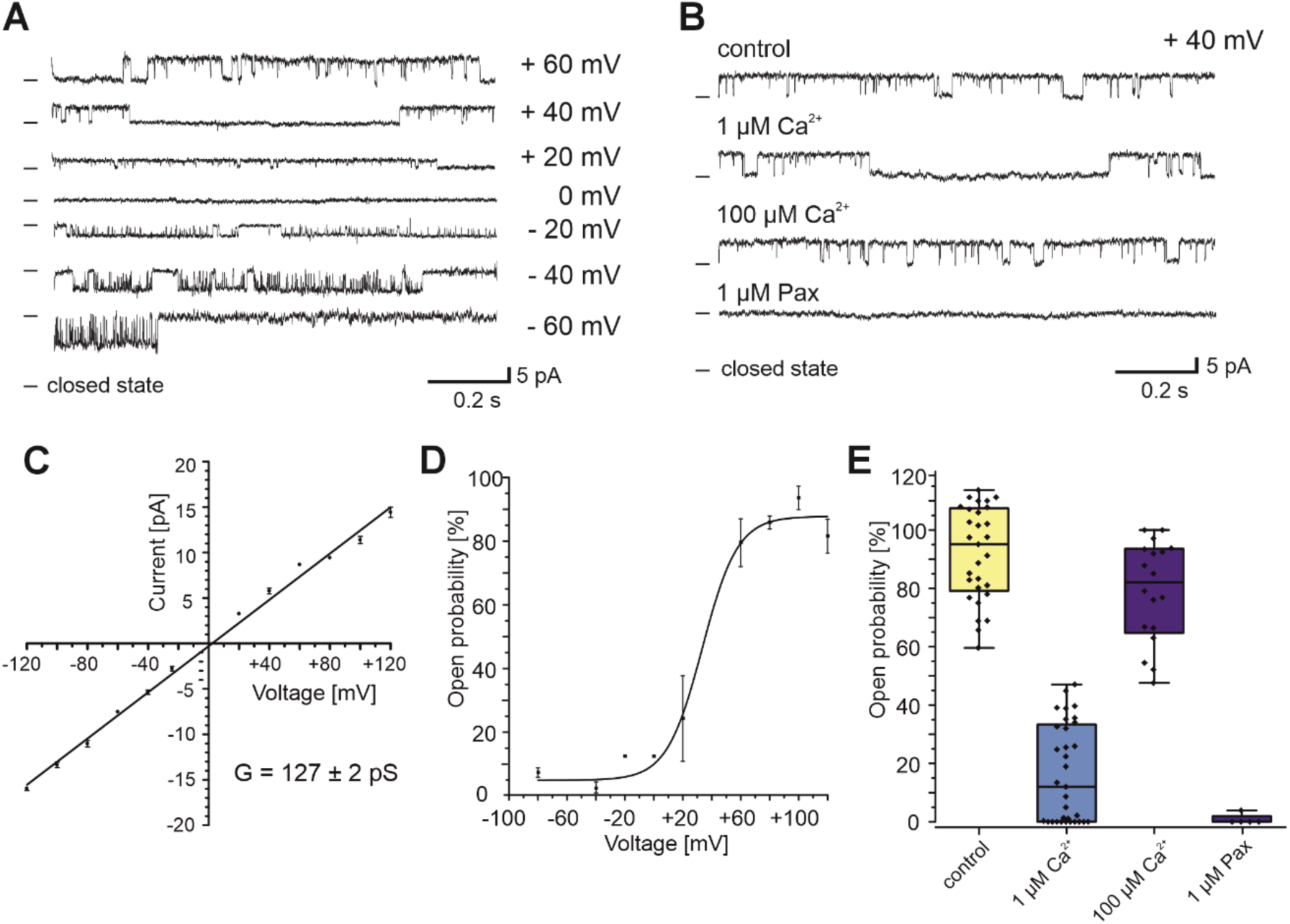
Electrophysiological characterization of mitochondrial large-conductance calcium-activated potassium (mitoBK_Ca_) channels in mitoplasts isolated from guinea pig cardiomyocytes. (A) Representative single-channel recordings obtained at different membrane potentials in symmetric 150/150 mM KCl isotonic solution in the presence of 100 µM Ca²⁺. (B) Representative single-channel current recordings measured at +40 mV under control conditions, in the presence of 1 µM and 100 µM Ca²⁺, after washout, and following application of 1 µM paxilline (Pax). In the panels A and B, the symbol “–” indicates the closed state of the channel. (C) Current–voltage (I–V) relationship of the mitoBK_Ca_ channels recorded in symmetric 150/150 mM KCl solution containing 100 µM Ca²⁺ (n = 4). The calculated single-channel conductance was equal to 127 ± 2 pS. (D) Open probability (Po) analysis in the presence of 100 µM Ca²⁺ analyzed over the voltage range from −80 to +120 mV (n = 4). (E) Quantitative analysis of mitoBK_Ca_ channel open probability under different experimental conditions: control, 1 µM Ca²⁺, 100 µM Ca²⁺, and 1 µM paxilline at +40 mV (n = 22). Data in panels B, C, and E are presented as mean ± SD.

**Figure 3.**
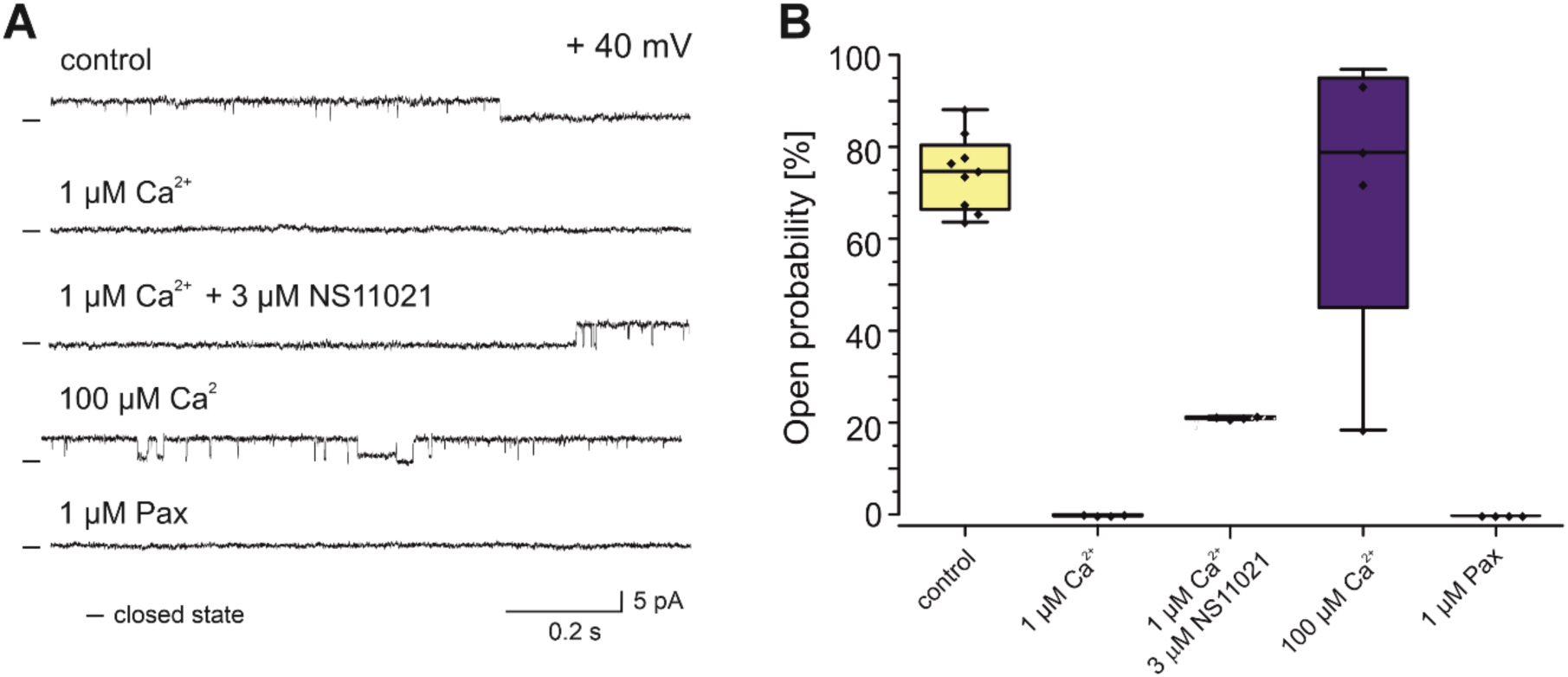
Electrophysiological characterization of mitoBK_Ca_ in the presence of the potassium channel opener NS11021. (A) Representative single-channel recordings obtained at +40 mV under control conditions, in the presence of 1 µM Ca²⁺, after application of 3 µM NS11021 in the presence of 1 µM Ca²⁺, in the presence of 100 µM Ca²⁺, and after addition of 1 µM paxilline. Application of 1 µM Ca²⁺ reduced channel activity, whereas NS11021 restored channel openings. Increasing Ca²⁺ concentration to 100 µM markedly enhanced channel activity. Paxilline almost completely inhibited channel openings. (B) Quantitative analysis of channel open probability under the indicated experimental conditions. Open probability was high under control conditions, decreased in the presence of 1 µM Ca²⁺, partially restored upon addition of NS11021, increased at 100 µM Ca²⁺, and was nearly abolished by 1 µM Paxilline. These results confirm calcium-dependent regulation of mitoBK_Ca_ channels and their pharmacological modulation by NS11021 and Paxilline (n = 7).

### Regulation of the mitoBK_Ca_ channel by infrared light

To investigate the regulation of the mitoBK_Ca_ channel by infrared light, we used the patch-clamp technique with an 820 nm illuminator. The results were summarized in Figure 4. Figure 4A shows plots of the channel open probability calculated from control conditions with 100 μM Ca^2+^; the channel exhibits relatively high activity. Addition of 300 μM K_3_[Fe(CN)_6_] strongly suppresses channel opening, reducing the open probability almost to zero. Exposure to 820 nm infrared light markedly restores channel activity despite the presence of K_3_[Fe(CN)_6_]. When the light is turned off, channel activity decreases again. Application of 1 μM paxilline (Pax), a selective BK_Ca_ channel inhibitor, nearly completely abolishes channel activity. Statistical significance is indicated by asterisks. Figure 4B presents representative single-channel current recordings corresponding to the experimental conditions shown in Figure 4A. In control conditions, frequent channel openings are observed. K_3_[Fe(CN)_6_] induces a mostly closed state, whereas simultaneous illumination with 820 nm light restores channel openings. Turning the 820 nm light off reduces activity again, and paxilline fully blocks the channel.

**Figure 4.**
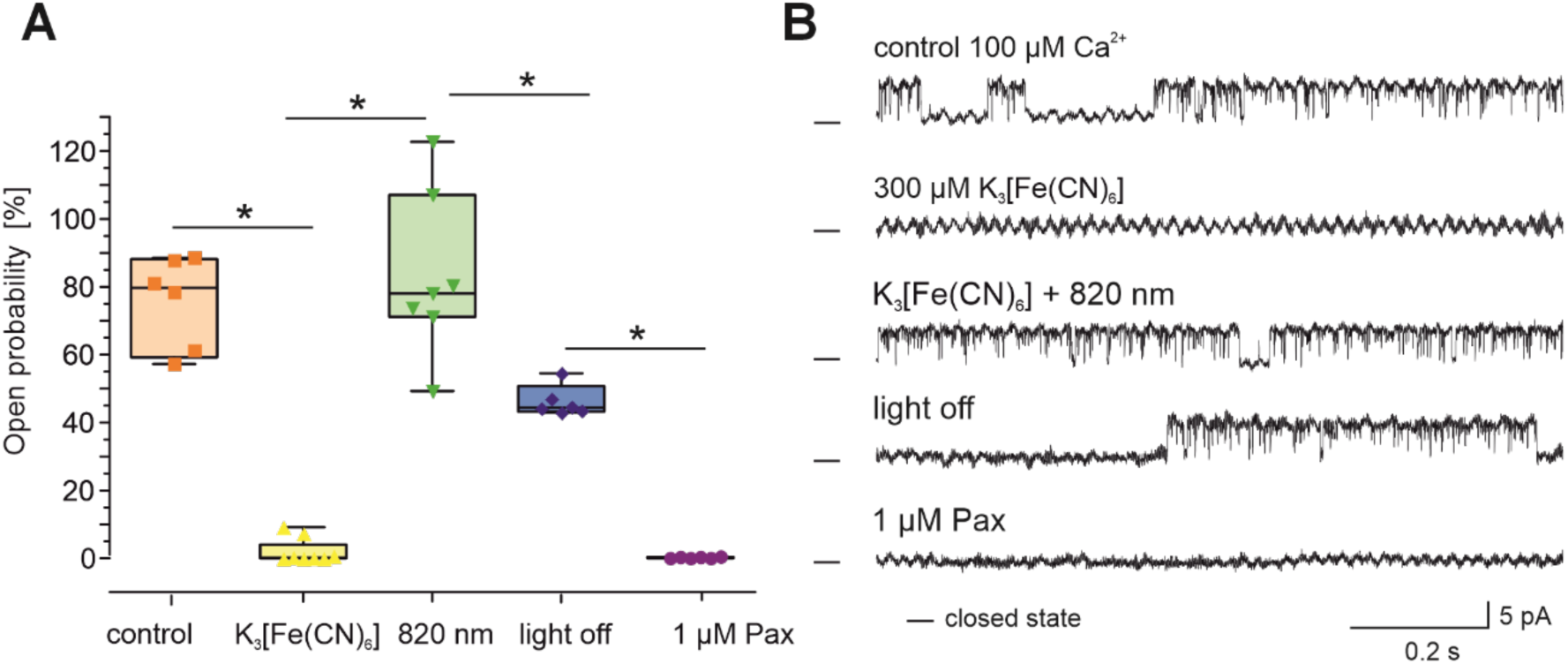
Regulation of the mitoBK_Ca_ channel activity by 820 nm infrared light. (A) plots showing the open probability of the mitoBK_Ca_ channel under different experimental conditions. In control conditions (100 μM Ca^2+^), the channel displayed high activity. The addition of 300 μM K_3_[Fe(CN)_6_] markedly reduced the channel open probability. Illumination with 820 nm infrared light restored channel activity in the presence of K_3_[Fe(CN)_6_]. After the light was switched off, channel activity decreased again. Application of 1 μM paxilline (Pax) almost completely inhibited channel openings. Data are presented with statistical significance indicated by * (n = 6). (B) Representative single-channel current traces recorded under the conditions shown in panel A. Infrared light increased mitoBK_Ca_ channel activity, suppressed by K_3_[Fe(CN)_6_], whereas paxilline fully blocked channel openings.

### Effect of 820 nm infrared light on hypoxic cardiomyocytes viability

In the final part of the study, we aimed to determine whether illumination with NIR light, which activates the mitoBK_Ca_ channel, could increase the survival of cardiomyocytes subjected to hypoxia. The obtained results indicate that hypoxic conditions significantly reduce cardiomyocyte viability compared with normoxia, confirming the damaging effect of hypoxia on cell survival. Under normoxic conditions, exposure to 820 nm infrared radiation increased cell viability; however, this effect did not reach statistical significance, suggesting only a tendency toward a protective effect under physiological conditions. Under hypoxic conditions, 820 nm infrared radiation statistically significantly attenuated the decline in cardiomyocyte viability (Figure 5). Importantly, the level of cell survival in this group was no longer significantly different from that observed in the control group under normoxic conditions, indicating a pronounced cytoprotective effect of 820 nm light under hypoxic stress. In summary, 820 nm irradiation has a limited effect under normoxic conditions but exerts a protective effect under hypoxic conditions, significantly improving cardiomyocyte viability and partially counteracting the adverse effects of hypoxia.

**Figure 5.**
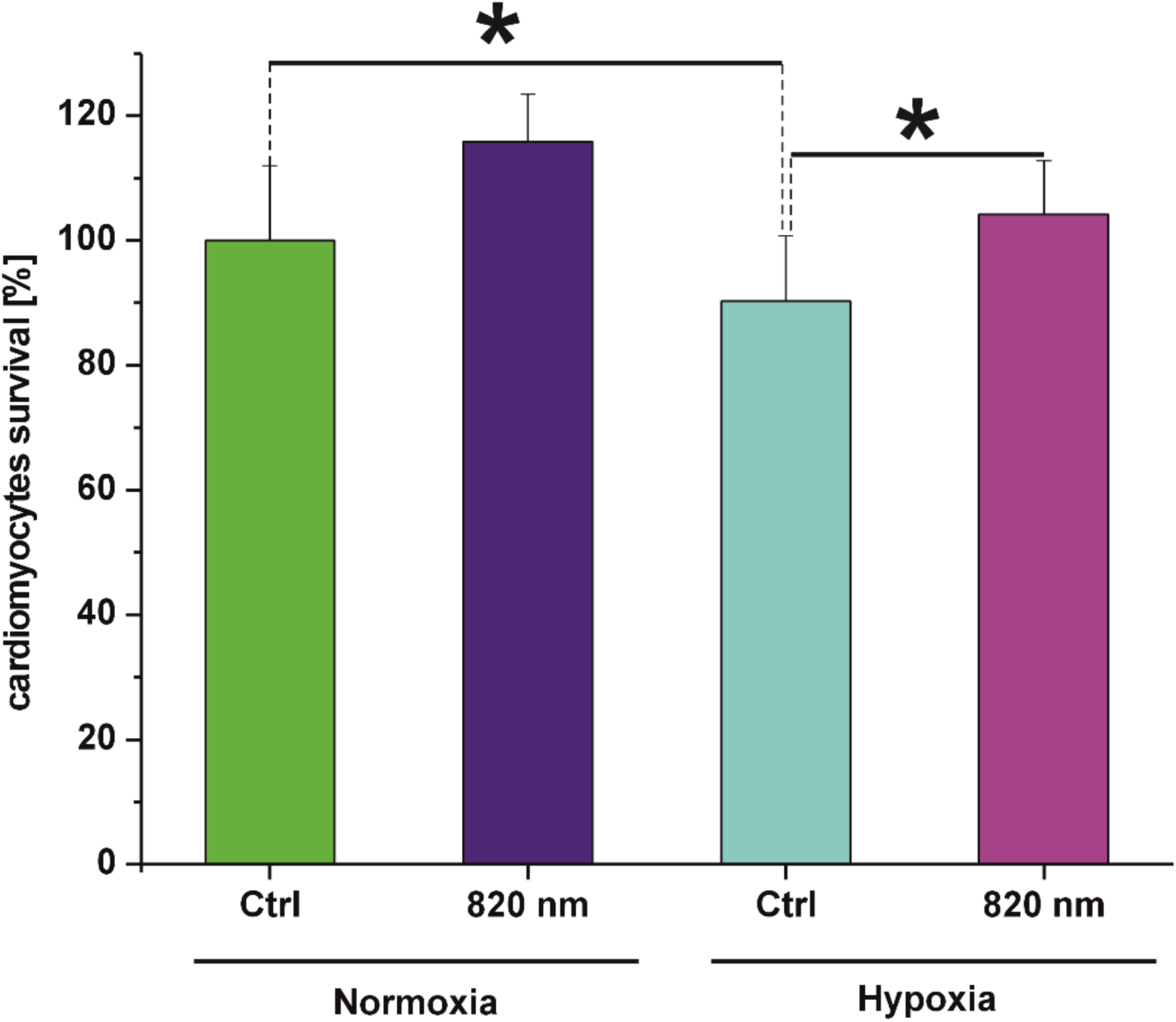
Effect of 820 nm infrared light on cardiomyocytes isolated from guinea pig hearts subjected to hypoxia. Cardiomyocyte viability was expressed as a percentage of the control value (100%) under both normoxic and hypoxic conditions. Under normoxia, exposure to 820 nm irradiation resulted in a non-significant increase in cell viability compared with the normoxic control group. Under hypoxia, viability in the control group was significantly reduced relative to the normoxic control. Illumination with 820 nm light partially attenuated this decrease, such that viability was no longer significantly different from the normoxic control group. Data are presented as mean ± SEM. Asterisks (*) denote statistically significant differences between groups (p < 0.05; n = 4).

**Figure 6.**
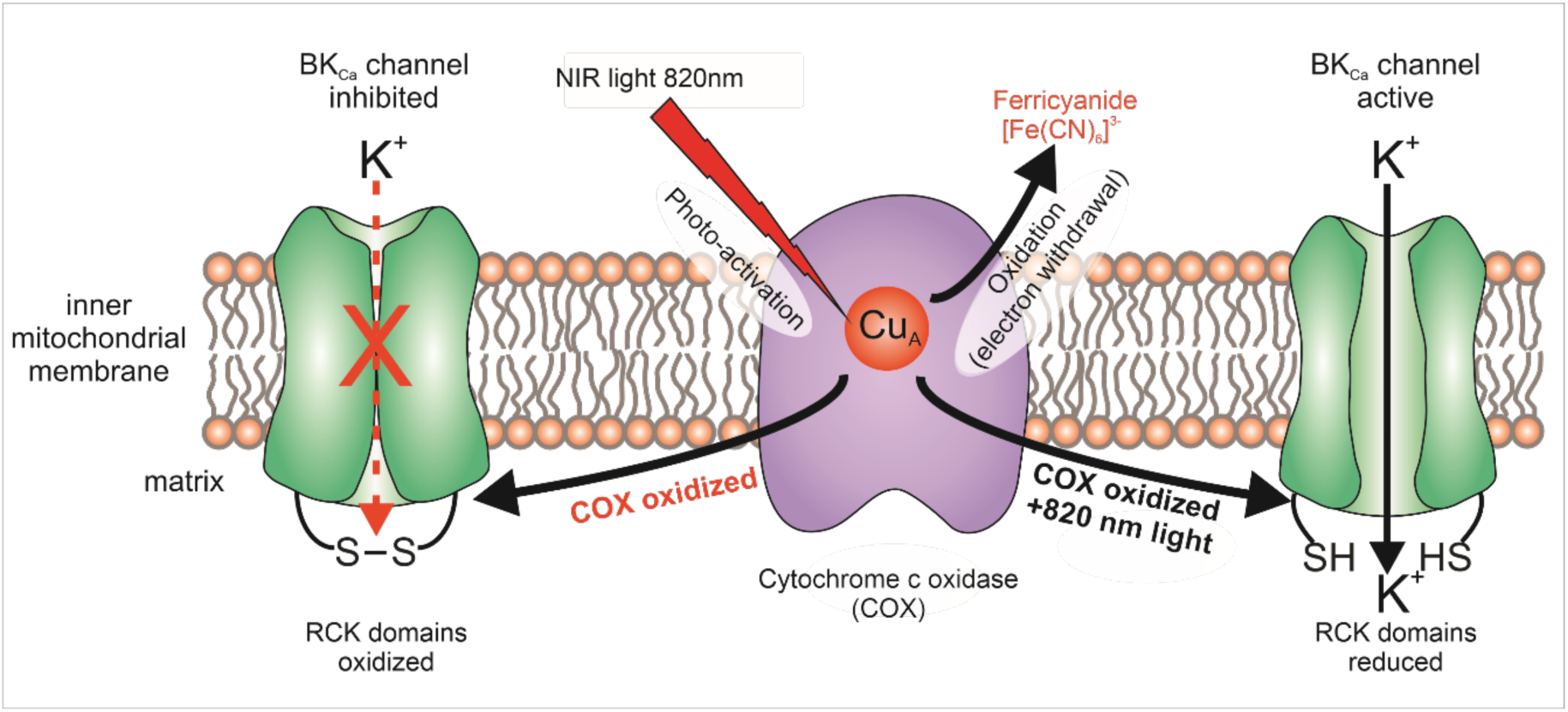
Proposed mechanism of mitoBK_Ca_ channel redox modulation via the COX Cu_A_ center. NIR light at 820 nm is selectively absorbed by the oxidized Cu_A_ center of cytochrome c oxidase (COX), accelerating intra-protein electron transfer and generating a local reductive environment. This process counteracts the oxidative inhibition caused by ferricyanide, which promotes disulfide bond (S-S) formation within the BK_Ca_ channel’s RCK domains. Although the BK_Ca_ channel lacks classical PAS domains, the RCK ring acts as a direct ‘redox switch’. The persistent activity observed after light cessation (hysteresis) reflects the slow kinetics of re-oxidation of the cysteine thiols (-SH) back to the inhibited S-S state.

## DISCUSSION

The results of the present study provide the first direct electrophysiological evidence that 820 nm infrared light stimulates the mitochondrial large-conductance calcium-activated potassium (mitoBK_Ca_) channel in guinea pig cardiomyocytes. Using patch-clamp recordings on isolated mitoplasts from cardiomyocytes, we have demonstrated that channel activation is intrinsically linked to the redox state of the mitochondrial respiratory chain and is likely modulated by the structural-functional coupling between the potassium channel and cytochrome c oxidase (COX). Our previous findings demonstrated that the activity of the astrocytoma mitoBK_Ca_ channel is modulated by the redox state of the respiratory chain (Bednarczyk et al., 2013).

### The cardiac mitoBK_Ca_ channel and the ‘Redox Window’

Our electrophysiological characterization confirms that the recorded mitoBK_Ca_ channel located in guinea pig cardiomyocytes exhibits typical BK-type channel properties, including high conductance and strong sensitivity to Ca^2+^, potential, and the specific potassium channel opener NS11021, which is consistent with previous reports (Bischof et al., 2024; Frankenreiter et al., 2017). However, the most significant finding concerns the modulation of mitoBK_Ca_ channels’ activity by the redox environment. As demonstrated, the application of potassium ferricyanide (K_3_[Fe(CN)_6_]) - a potent oxidizing agent - nearly abolished the channel open probability. Potassium ferricyanide acts as a potent electron acceptor directly from the respiratory chain components, such as cytochrome c or COX, thereby maintaining the mitochondrial electron transport system in a highly oxidized state. This observation aligns perfectly with the ‘redox window’ hypothesis (DiChiara & Reinhart, 1997), which postulates that BK channels operate optimally only within a specific, physiological redox range, whereas extreme oxidation shifts the system beyond this functional window. We previously observed a similar phenomenon with the mitoBK_Ca_ channel in astrocytoma cells, suggesting the existence of a conserved regulatory mechanism across tissues (Bednarczyk et al., 2026).

In our cardiac model, 820 nm infrared light acts as a “redox-reversing” stimulus. When the channel is inhibited by oxidative stress, illumination at 820 nm, a wavelength specifically absorbed by the Cu_A_ centers of COX, restores the mitoBK_Ca_ channel’s open probability. COX is considered the main mitochondrial chromophore for red and near-infrared light (600-850 nm) due to its heme a, heme a3/Cu_B_, and Cu_A_/Cu_B_ centers that undergo reversible redox transitions upon photon absorption (Dyuba et al., 2013; Farver et al., 2006; Mason et al., 2009, 2014; Szundi et al., 2001). This suggests that photon absorption by COX does not merely stimulate electron transport but actively shifts the local redox environment of the mitoBK_Ca_ channel back into its optimal functional ‘window’, effectively neutralizing the inhibitory effect of the applied oxidant, i.e., ferricyanide.

### Topological and structural coupling: the mitochondrial interactome

The rapid nature of the channel’s response to 820 nm light may support the existence of a cardiac mitochondrial ‘interactome’ (Caudal et al., 2022; Schweppe et al., 2017). While the physical distance between COX copper centers and the channel’s redox-sensitive cysteines might theoretically limit signal transfer, evidence suggests a close spatial association. In cardiomyocytes, the β1 regulatory subunit of the mitoBK_Ca_ channel interacts with the COX subunit I (Ohya et al., 2005), while studies in other models have confirmed functional interactions between the channel and the respiratory chain (Bednarczyk et al., 2013). This probable structural tethering drastically reduces the spatial barrier, allowing for near-instantaneous transmission of light-induced redox shifts from COX to the channel’s gating machinery via conformational signaling and changes in redox-sensitive sites of the mitoBK_Ca_ channel.

### Chemical-conformational “Memory” and hysteresis

A striking feature of our recordings of mitoBK_Ca_ channel activity from cardiomyocyte mitochondria is the “memory” effect observed after 820 nm illumination was discontinued. Turning off the light decreased channel activity, but the channel open probability remained significantly higher than the baseline pre-illumination oxidation level. This hysteresis suggests that 820 nm light induces stable chemical-conformational changes in the mitoBK_Ca_ protein. Such durability indicates that the underlying mechanism is not based on a transient photothermal effect, which would dissipate rapidly. Instead, it involves modulation of a redox sensor whose relaxation time is limited by the slow kinetics of chemical re-oxidation within the local COX – mitoBK_Ca_ channel microenvironment. This maintains the channel within its optimal ‘redox window’ even after the infra-red light stimulus is eliminated.

Our electrophysiological observations suggest a functional coupling between COX and the mitoBK_Ca_ channel, mediated by a redox-dependent mechanism rather than direct photoactivation. The inhibition of mitoBK_Ca_ channel activity by ferricyanide, a potent oxidizing agent, underscores the channel’s sensitivity to the redox state, likely through the oxidation of critical cysteine residues within the C-terminal RCK (Regulator of Conductance for Potassium) domains.

Conversely, the application of 820 nm light, which specifically matches the absorption band of the oxidized Cu_A_ center in COX, significantly stimulated channel activity. We propose that the absorption of near-infrared (NIR) photons by the Cu_A_ center effectively shifts the local microenvironment toward a more reduced state. This light-induced effect may also facilitate the dissociation of inhibitory molecules, such as nitric oxide (NO), or generate transient redox signals that act on the BK_Ca_ channel’s RCK sensors, lowering the energy barrier for pore opening (Hamblin, 2018).

While many oxygen and redox-sensitive proteins rely on specialized PAS (Per-Arnt-Sim) domains to transduce environmental signals (Dhungel et al., 2025; Xing et al., 2023), our results reinforce the conclusion that the mitoBK_Ca_ channel utilizes a distinct, more direct sensory architecture. We hypothesize that the mitoBK_Ca_ channel’s redox sensitivity is achieved through the integration of its RCK (Regulator of Conductance for Potassium) domains with intrinsic cytochrome-like motifs (such as the C-terminal heme-binding or CKACH sequences) (Gudratova et al., 2026; Yusifov et al., 2025). This evolutionary adaptation bypasses the need for classical PAS domains, allowing the channel to function as a primary redox-coupled pore that responds instantaneously to redox transitions within COX. By operating as a direct ‘redox-to-conductance’ transducer, the mitoBK_Ca_ channel ensures rapid synchronization of mitochondrial membrane potential with the cell’s metabolic rate. This could be followed by the level of ROS synthesis by intact mitochondria (Szabo and Szewczyk, 2023).

An interesting finding in our study is the ‘hysteresis’ effect: upon stopping 820 nm illumination, the channel activity declined but remained above baseline levels. This persistent activation suggests that NIR light triggers downstream biochemical modifications, such as the reduction of disulfide bonds, which do not immediately revert upon removal of the 820 nm light stimulus. This ‘redox memory’ further supports the hypothesis that the mitoBK_Ca_ channel serves as a downstream effector of COX-mediated mitochondrial signaling. Notably, as the mitoBK_Ca_ channel lacks classical PAS domains, this sensitivity is likely conferred by its intrinsic redox-sensitive motifs or by its direct physical association with the respiratory complex, allowing a synchronized response to metabolic and infrared light stimuli.

Our findings demonstrate that the mitoBK_Ca_ channel acts as a metabolic sensor, finely tuned to the redox status of the mitochondrial respiratory chain. The stimulation of channel activity by 820 nm light, coupled with the inhibitory effect of ferricyanide, confirms that the mitoBK_Ca_ channel’s gating is dynamically regulated by the oxidation state of cytochrome c oxidase. The observed persistence of channel activation post-illumination suggests that NIR light induces relatively stable covalent changes (e.g., thiol-disulfide exchange) within the mitoBK_Ca_ channel’s regulatory apparatus, providing a mechanism for sustained cytoprotection following photobiomodulation.

The observed functional synchrony between COX and mitoBK_Ca_ channel gating may be further facilitated by the presence of accessory β subunits. While the α subunit contains the core RCK domains responsible for redox sensing, evidence suggests that β subunits, particularly the β1, physically and functionally associate with mitochondrial respiratory complexes, including COX (Ohya et al., 2005). In our model, we hypothesize that the β subunit may act as a structural transducer, bridging the gap between the COX Cu_A_ center and the mitoBK_Ca_ channel’s gating ring. Such an interaction could optimize the transmission of the light-induced reductive signal, possibly by modulating the conformational accessibility of key cysteine residues within the RCK domains. Thus, while the mitoBK_Ca_ channel lacks classical PAS domains, the inclusion of β subunits likely amplifies its sensitivity to NIR-induced redox shifts, ensuring a rapid response to changes in mitochondrial metabolic status.

### Cardioprotective implications of infrared illumination

The physiological significance of NIR light mitoBK_Ca_ channels regulation is highlighted by our survival assays. Preconditioning with 820 nm light significantly increased the viability of hypoxic cardiomyocytes, matching the survival rates of normoxic controls. This protective effect is likely mediated by light-induced opening of mitoBK_Ca_ channels, which facilitates K^+^ influx and modulates mitochondrial function. These findings identify 820 nm light as a potent non-pharmacological tool for maintaining mitochondrial homeostasis and enhancing cardiac resistance against ischemia/reperfusion stress.

### Final remarks

Our results demonstrate that the mitoBK_Ca_ channel functions as a metabolic sensor functionally coupled to the mitochondrial respiratory chain. We propose that NIR light within COX triggers a local redox shift that is transduced to the mitoBK_Ca_ channel, probably via its RCK (Regulator of Conductance for Potassium) domains. This direct coupling bypasses the need for PAS domains, allowing for rapid, sustained synchronization between mitochondrial respiration and K^+^ ion conductance. These findings provide a new molecular basis for the cytoprotective effects observed in photobiomodulation therapy.

## ACKNOWLEDGMENTS

Supported by the Polish National Science Center (MAESTRO grant No. 2019/34/A/NZ1/00352).

## REFERENCES

1. Abijo, A., Lee, C. Y., Huang, C. Y., Ho, P. C., & Tsai, K. J. (2023). The Beneficial Role of Photobiomodulation in Neurodegenerative Diseases. Biomedicines, 11(7). 10.3390/BIOMEDICINES11071828

2. Ardehali, H., Chen, Z., Ko, Y., Mejía-Alvarez, R., & Marbán, E. (2004). Multiprotein complex containing succinate dehydrogenase confers mitochondrial ATP-sensitive K^+^ channel activity. Proceedings of the National Academy of Sciences of the United States of America, 101(32), 11880–11885. 10.1073/PNAS.0401703101

3. Augustynek, B., Kunz, W. S., & Szewczyk, A. (2017). Guide to the Pharmacology of Mitochondrial Potassium Channels. Handbook of Experimental Pharmacology, 240. 10.1007/164_2016_79

4. Bednarczyk, P., Lewandowska, J., Kulawiak, B., & Szewczyk, A. (2026). Red/near-infrared light activates the mitochondrial large-conductance calcium-activated potassium channel in glioblastoma cells. BioRxiv, 2026.04.02.716077. 10.64898/2026.04.02.716077

5. Bednarczyk, P., Wieckowski, M. R., Broszkiewicz, M., Skowronek, K., Siemen, D., & Szewczyk, A. (2013). Putative Structural and Functional Coupling of the Mitochondrial BKCa Channel to the Respiratory Chain. PloS One, 8(6). 10.1371/JOURNAL.PONE.0068125

6. Bischof, H., Maier, S., Koprowski, P., Kulawiak, B., Burgstaller, S., Jasińska, J., Serafimov, K., Zochowska, M., Gross, D., Schroth, W., Matt, L., Lopez, D. A. J., Zhang, Y., Bonzheim, I., Büttner, F. A., Fend, F., Schwab, M., Birkenfeld, A. L., Malli, R., … Lukowski, R. (2024). mitoBKCa is functionally expressed in murine and human breast cancer cells and potentially contributes to metabolic reprogramming. ELife, 12. 10.7554/ELIFE.92511.2

7. Caudal, A., Tang, X., Chavez, J. D., Keller, A., Mohr, J. P., Bakhtina, A. A., Villet, O., Chen, H., Zhou, B., Walker, M. A., Tian, R., & Bruce, J. E. (2022). Mitochondrial interactome quantitation reveals structural changes in metabolic machinery in the failing murine heart. Nature Cardiovascular Research 1(9), 855–866. 10.1038/s44161-022-00127-4

8. Chandel, N. S. (2014). Mitochondria as signaling organelles. BMC Biology, 12(1), 34. 10.1186/1741-7007-12-34

9. Dhungel, S., Xiao, M., Pushpabai, R. R., & Kikani, C. K. (2025). Structural assembly of the PAS domain drives the catalytic activation of metazoan PASK. Proceedings of the National Academy of Sciences of the United States of America, 122(12), e2409685122. 10.1073/PNAS.2409685122

10. DiChiara, T. J., & Reinhart, P. H. (1997). Redox modulation of *hslo* Ca^2+^-activated K^+^ channels. The Journal of Neuroscience: The Official Journal of the Society for Neuroscience, 17(13), 4942–4955. 10.1523/JNEUROSCI.17-13-04942.1997

11. Dyuba, A. V., Vygodina, T. V., & Konstantinov, A. A. (2013). Reconstruction of absolute absorption spectrum of reduced heme a in cytochrome C oxidase from bovine heart. Biochemistry. Biokhimiia, 78(12), 1358–1365. 10.1134/S0006297913120067

12. Farver, O., Grell, E., Ludwig, B., Michel, H., & Pecht, I. (2006). Rates and equilibrium of CuA to heme a electron transfer in Paracoccus denitrificans cytochrome c oxidase. Biophysical Journal, 90(6), 2131–2137. 10.1529/biophysj.105.075440

13. Frankenreiter, S., Bednarczyk, P., Kniess, A., Bork, N. I., Straubinger, J., Koprowski, P., Wrzosek, A., Mohr, E., Logan, A., Murphy, M. P., Gawaz, M., Krieg, T., Szewczyk, A., Nikolaev, V. O., Ruth, P., & Lukowski, R. (2017). cGMP-Elevating Compounds and Ischemic Conditioning Provide Cardioprotection Against Ischemia and Reperfusion Injury via Cardiomyocyte-Specific BK Channels. Circulation, 136(24), 2337–2355. 10.1161/CIRCULATIONAHA.117.028723

14. Galluzzi, L., Kepp, O., & Kroemer, G. (2012). Mitochondria: master regulators of danger signalling. Nature Reviews. Molecular Cell Biology, 13(12), 780–788. 10.1038/NRM3479

15. Garlid, K. D., Dos Santos, P., Xie, Z. J., Costa, A. D. T., & Paucek, P. (2003). Mitochondrial potassium transport: The role of the mitochondrial ATP-sensitive K^+^ channel in cardiac function and cardioprotection. Biochimica et Biophysica Acta - Bioenergetics, 1606(1–3), 1–21. 10.1016/S0005-2728(03)00109-9

16. Gorman, A. M., Healy, S. J. M., Jäger, R., & Samali, A. (2012). Stress management at the ER: Regulators of ER stress-induced apoptosis. Pharmacology and Therapeutics, 134(3), 306–316. 10.1016/j.pharmthera.2012.02.003

17. Gudratova, F., Savalli, N., Aliyeva, A., & Yusifov, T. (2026). Heme signaling through the cytochrome c-like domain of the human BK channel. Biochimica et Biophysica Acta - Biomembranes, 1868(3), 184533. 10.1016/j.bbamem.2026.184533

18. Hamblin, M. R. (2018). Mechanisms and Mitochondrial Redox Signaling in Photobiomodulation. Photochemistry and Photobiology, 94(2), 199–212. 10.1111/PHP.12864

19. Kampa, R. P., Sęk, A., Szewczyk, A., & Bednarczyk, P. (2021). Cytoprotective effects of the flavonoid quercetin by activating mitochondrial BKCa channels in endothelial cells. Biomedicine and Pharmacotherapy, 142. 10.1016/j.biopha.2021.112039

20. Karu, T. I. (2010). Multiple roles of cytochrome c oxidase in mammalian cells under action of red and IR-A radiation. IUBMB Life, 62(8), 607–610. 10.1002/IUB.359

21. Karu, T. I., Pyatibrat, L. V., & Kalendo, G. S. (2004). Photobiological modulation of cell attachment via cytochrome c oxidase. Photochemical and Photobiological Sciences, 3(2), 211–216. 10.1039/b306126d

22. Kathiresan, T., Harvey, M., Orchard, S., Sakai, Y., & Sokolowski, B. (2009). A protein interaction network for the large conductance Ca ^2+^-activated K^+^ channel in the mouse cochlea. Molecular and Cellular Proteomics, 8(8), 1972–1987. 10.1074/mcp.M800495-MCP200

23. Kwong, J. Q., Lu, X., Correll, R. N., Schwanekamp, J. A., Vagnozzi, R. J., Sargent, M. A., York, A. J., Zhang, J., Bers, D. M., & Molkentin, J. D. (2015). The Mitochondrial Calcium Uniporter Selectively Matches Metabolic Output to Acute Contractile Stress in the Heart. Cell Reports, 12(1), 15–22. 10.1016/j.celrep.2015.06.002

24. Lewandowska, J., Kalenik, B., Szewczyk, A., & Wrzosek, A. (2026). Rapid protocol for mitochondria isolation from cardiomyocytes employing cell strainer-based procedure. BioRxiv, 2026.04.02.716092. 10.64898/2026.04.02.716092

25. Liu, Y., Gao, W. D., O’Rourke, B., & Marban, E. (1996). Cell-type specificity of preconditioning in an in vitro model. Basic Research in Cardiology, 91(6), 450–457. 10.1007/BF00788726

26. Mason, M. G., Nicholls, P., & Cooper, C. E. (2009). The steady-state mechanism of cytochrome c oxidase: redox interactions between metal centres. The Biochemical Journal, 422(2), 237–246. 10.1042/BJ20082220

27. Mason, M. G., Nicholls, P., & Cooper, C. E. (2014). Re-evaluation of the near infrared spectra of mitochondrial cytochrome c oxidase: Implications for non invasive in vivo monitoring of tissues. Biochimica et Biophysica Acta - Bioenergetics, 1837(11), 1882–1891. 10.1016/j.bbabio.2014.08.005

28. Murphy, M. P. (2009). How mitochondria produce reactive oxygen species. The Biochemical Journal, 417(1), 1–13. 10.1042/BJ20081386

29. Ohya, S., Kuwata, Y., Sakamoto, K., Muraki, K., & Imaizumi, Y. (2005). Cardioprotective effects of estradiol include the activation of large-conductance Ca^2+^-activated K^+^ channels in cardiac mitochondria. American Journal of Physiology. Heart and Circulatory Physiology, 289(4), H1635–H1642. 10.1152/AJPHEART.00016.2005

30. O’Rourke, B. (2004). Evidence for mitochondrial K^+^ channels and their role in cardioprotection. Circulation Research, 94(4), 420–432. 10.1161/01.RES.0000117583.66950.43

31. O’Rourke, B. (2007). Mitochondrial ion channels. Annual Review of Physiology, 69, 19–49. 10.1146/ANNUREV.PHYSIOL.69.031905.163804

32. Peng, Z., Sakai, Y., Kurgan, L., Sokolowski, B., & Uversky, V. (2014). Intrinsic disorder in the BK channel and its interactome. PloS One, 9(4), e94331. 10.1371/JOURNAL.PONE.0094331

33. Peruzzo, R., Mattarei, A., Azzolini, M., Becker-Flegler, K. A., Romio, M., Rigoni, G., Carrer, A., Biasutto, L., Parrasia, S., Kadow, S., Managò, A., Urbani, A., Rossa, A., Semenzato, G., Soriano, M. E., Trentin, L., Ahmad, S., Edwards, M., Gulbins, E., … Szabò, I. (2020). Insight into the mechanism of cytotoxicity of membrane-permeant psoralenic Kv1.3 channel inhibitors by chemical dissection of a novel member of the family. Redox Biology, 37, 101705. 10.1016/j.redox.2020.101705

34. Pham, L., Arroum, T., Morse, P. T., Bell, J., Malek, M. H., Sanderson, T. H., & Hüttemann, M. (2025). Inhibitory Infrared Light Attenuates Mitochondrial Hyperactivity and Accelerates Restoration of Mitochondrial Homeostasis in an Oxygen-Glucose Deprivation/Reoxygenation Model. Antioxidants (Basel, Switzerland), 14(9), 1119. 10.3390/ANTIOX14091119

35. Sanderson, T. H., Wider, J. M., Lee, I., Reynolds, C. A., Liu, J., Lepore, B., Tousignant, R., Bukowski, M. J., Johnston, H., Fite, A., Raghunayakula, S., Kamholz, J., Grossman, L. I., Przyklenk, K., & Hüttemann, M. (2018). Inhibitory modulation of cytochrome c oxidase activity with specific near-infrared light wavelengths attenuates brain ischemia/reperfusion injury. Scientific Reports, 8(1), 3481. 10.1038/s41598-018-21869-x

36. Schweppe, D. K., Chavez, J. D., Lee, C. F., Caudal, A., Kruse, S. E., Stuppard, R., Marcinek, D. J., Shadel, G. S., Tian, R., & Bruce, J. E. (2017). Mitochondrial protein interactome elucidated by chemical cross-linking mass spectrometry. Proceedings of the National Academy of Sciences of the United States of America, 114(7), 1732–1737. 10.1073/PNAS.1617220114

37. Siemen, D., Loupatatzis, C., Borecky, J., Gulbins, E., & Lang, F. (1999). Ca^2+^-activated K channel of the BK-type in the inner mitochondrial membrane of a human glioma cell line. Biochemical and Biophysical Research Communications, 257(2), 549–554. 10.1006/BBRC.1999.0496

38. Singh, H., Li, M., Hall, L., Chen, S., Sukur, S., Lu, R., Caputo, A., Meredith, A. L., Stefani, E., & Toro, L. (2016). MaxiK channel interactome reveals its interaction with GABA transporter 3 and heat shock protein 60 in the mammalian brain. Neuroscience, 317, 76–107. 10.1016/j.neuroscience.2015.12.058

39. Sokolowski, B., Orchard, S., Harvey, M., Sridhar, S., & Sakai, Y. (2011). Conserved BK channel-protein interactions reveal signals relevant to cell death and survival. PloS One, 6(12), e28532. 10.1371/JOURNAL.PONE.0028532

40. Szabo, I., & Szewczyk, A. (2023). Mitochondrial Ion Channels. Annual Review of Biophysics, 52, 229–254. 10.1146/ANNUREV-BIOPHYS-092622-094853

41. Szabo, I., & Zoratti, M. (2014). Mitochondrial channels: ion fluxes and more. Physiological Reviews, 94(2), 519–608. 10.1152/PHYSREV.00021.2013

42. Szteyn, K., & Singh, H. (2020). BK_Ca_ Channels as Targets for Cardioprotection. *Antioxidants (Basel*, Switzerland), 9(8), 1–16. 10.3390/ANTIOX9080760

43. Szundi, I., Liao, G. L., & Einarsdóttir, O. (2001). Near-infrared time-resolved optical absorption studies of the reaction of fully reduced cytochrome c oxidase with dioxygen. Biochemistry, 40(8), 2332–2339. 10.1021/BI002220V

44. Walewska, A., Krajewska, M., Stefanowska, A., Buta, A., Bilewicz, R., Krysiński, P., Bednarczyk, P., Koprowski, P., & Szewczyk, A. (2022). Methods of Measuring Mitochondrial Potassium Channels: A Critical Assessment. International Journal of Molecular Sciences, 23(3). 10.3390/IJMS23031210

45. Wider, J. M., Gruley, E., Morse, P. T., Wan, J., Lee, I., Anzell, A. R., Fogo, G. M., Mathieu, J., Hish, G., O’Neil, B., Neumar, R. W., Przyklenk, K., Hüttemann, M., & Sanderson, T. H. (2023). Modulation of mitochondrial function with near-infrared light reduces brain injury in a translational model of cardiac arrest. Critical Care (London, England), 27(1), 491, eadi4517. 10.1186/S13054-023-04745-7

46. Xing, J., Gumerov, V. M., & Zhulin, I. B. (2023). Origin and functional diversification of PAS domain, a ubiquitous intracellular sensor. Science Advances, 9(35), eadi4517. 10.1126/SCIADV.ADI4517

47. Xu, W., Liu, Y., Wang, S., McDonald, T., Van Eyk, J. E., Sidor, A., & O’Rourke, B. (2002). Cytoprotective role of Ca^2+^-activated K^+^ channels in the cardiac inner mitochondrial membrane. Science (New York, N.Y.), 298(5595), 1029–1033. 10.1126/SCIENCE.1074360

48. Yao, J., McHedlishvili, D., McIntire, W. E., Guagliardo, N. A., Erisir, A., Coburn, C. A., Santarelli, V. P., Bayliss, D. A., & Barrett, P. Q. (2017). Functional TASK-3-Like Channels in Mitochondria of Aldosterone-Producing Zona Glomerulosa Cells. Hypertension, 70(2), 347–356. 10.1161/HYPERTENSIONAHA.116.08871

49. Yusifov, T., Qudretova, F., & Aliyeva, A. (2025). Cytochrome C-like Domain Within the Human BK Channel. International Journal of Molecular Sciences, 26(15), 7053. 10.3390/IJMS26157053

50. Zhang, J., Li, M., Zhang, Z., Zhu, R., Olcese, R., Stefani, E., & Toro, L. (2017). The mitochondrial BK_Ca_ channel cardiac interactome reveals BK_Ca_ association with the mitochondrial import receptor subunit Tom22, and the adenine nucleotide translocator. Mitochondrion, 33, 84–101. 10.1016/j.mito.2016.08.017

51. Zhang, W., Gao, X., Wang, X., Li, D., Zhao, Y., Zhang, T., Ne, J., Xu, B., Li, S., Jiang, Z., Sun, H., Ma, W., Yang, F., Cai, B., & Yang, B. (2021). Light Emitting Diodes Photobiomodulation Improves Cardiac Function by Promoting ATP Synthesis in Mice With Heart Failure. Frontiers in Cardiovascular Medicine, 8, 753664. 10.3389/fcvm.2021.753664

